# Evolutionarily diverse fungal zoospores show contrasting swimming patterns specific to ultrastructure

**DOI:** 10.1101/2023.01.22.525074

**Authors:** Luis Javier Galindo, Thomas A. Richards, Jasmine A. Nirody

## Abstract

Zoosporic fungi, also called chytrids, produce motile spores with flagellar swimming tails (zoospores)^1,2^. These fungi are key components of aquatic food webs, acting as pathogens, saprotrophs and prey^3–8^. Little is known about the swimming behaviour of fungal zoospores, a crucial factor governing dispersal, biogeographical range, ecological function and infection dynamics^6,9^. Here, we track the swimming patterns of zoospores from 12 evolutionary divergent species of zoosporic fungi across seven orders of the Chytridiomycota and the Blastocladiomycota phyla. We report two major swimming patterns which correlate with the cytoskeletal ultrastructure of these zoospores. Specifically, we show that species without major cytoplasmic tubulin components swim in a circular fashion, while species that harbour prominent cytoplasmic tubulin structures swim in a pattern akin to a random walk (move-stop-redirect-move). We confirm cytoskeleton architecture by performing fluorescence confocal microscopy of the zoospores across all 12 species. We then treat representative species with variant swimming behaviours and cytoplasmic-cytoskeletal arrangements with tubulin stabilizing (Taxol) and depolymerizing (Nocodazole) pharmacological-compounds. We observed that when treating the ‘random-walk’ species with Nocodazole their swimming behaviour changes to a circular swimming pattern. Confocal imaging of the nocodazole-treated zoospores demonstrates these cells maintain flagellum tubulin structures but lack their characteristic cytoplasmatic tubulin arrangement. These data confirm that the capability of zoospores to perform ‘complex’ movements as a random walk is linked to the presence of prominent cytoplasmatic tubulin structures. We discuss the link between cytology, sensation, and swimming behaviour manifest in zoosporic fungi.

## Results

### Differential swimming in variant chytrids

Here we analysed swimming patterns of zoospores from 12 different species from the two major chytrid phyla: Chytridiomycota and Blastocladiomycota. These include eight Chytridiomycota species grouped into six orders: *Synchytrium microbalum* (Synchytriales), *Clydaea vesicula* (Lobulomycetales), *Chytriomyces hyalinus* (Chytridiales), *Rhizoclosmatium globosum* (Chytridiales), *Homolaphlyctis polyrhiza* (Rhizophydiales), *Rhizophlyctis rosea* (Rhizophlyctidiales), *Geranomyces variabilis* (Spizellomycetales), and *Spizellomyces punctatus* (Spizellomycetales); and four species of Blastocladiomycota from the order Blastocladiales: *Catenophlyctis* sp. (Catenariaceae), *Blastocladiella emersonii*, *Allomyces macrogynus* and *Allomyces reticulatus* (Blastocladiaceae).

We find that medium-range motility patterns varied markedly among different species (Figure 1), with the zoospores from species of four Chytridiomycota (e.g., *S. microbalum*, *C. hyalinus*, *R. globosum* and *H. polyrhiza*) swimming in a circular mode. In contrast, all zoospores from species of Blastocladiomycota tested and four different species of Chytridiomycota (e.g., *C. vesicula*, *R. rosea*, *G. variabilis* and *S. punctatus*) swim using a pattern akin to a random walk (move-stop-redirect-move; Figure 1A). To investigate this pattern further, we plotted the distribution of reorientation (turn) angles and the mean square displacement (MSD) of these 12 movement patterns to characterize motility dynamics at both shorter and longer timescales, respectively (see the methods section for the mathematical formulae). Such methods can be used to quantitatively differentiate random walk and circular motility patterns^10,11^. Circular swimming cells are expected to reorient themselves more consistently than those swimming in a random walk-like pattern, resulting in a narrower turn angle distribution. For cells exhibiting circular motility patterns, the MSD plot should form a plateau in the later phases of the MSDs plots. Consistent with the conclusions from our primary observations (Figure 1A), we find that reorientation angle distributions are indeed generally tighter for zoosporic fungi without prominent tubulin cytoplasmic structures (Figure 1B), and that the MSD plateaus at shorter timescales for these species in comparison to the other zoosporic fungi (Figure 1C). We note that the random-walk swimmer *A. macrogynus* (*Am*) shows a narrower distribution and the circular swimmer *C. hyalinus* (*Ch*) shows a broader angle range than expected. This is because *Am* tends to swim in more regular, directed trajectories than other random-walkers; this is also apparent from its MSD which scales with a higher exponent than that of other species (the inconsistencies of this species are discussed further below). On the other hand, *Ch* is the circular-swimmer with the most variable radii, resulting in a wider turn angle distribution. We expect that differences in local behaviours will be further heightened in experiments taken at greater resolution (e.g., using a camera with a higher frame rate). These results provide support for a primary conclusion that zoosporic fungi show two distinct swimming patterns. Apart from slightly higher speeds observed in *H. polyrhiza*, no significant differences in instantaneous swimming speed were observed across the 12 species (Figure S3A), suggesting the difference in movement behaviour is a product of cellular mechanics and not propulsion velocity.

**Figure 1.**
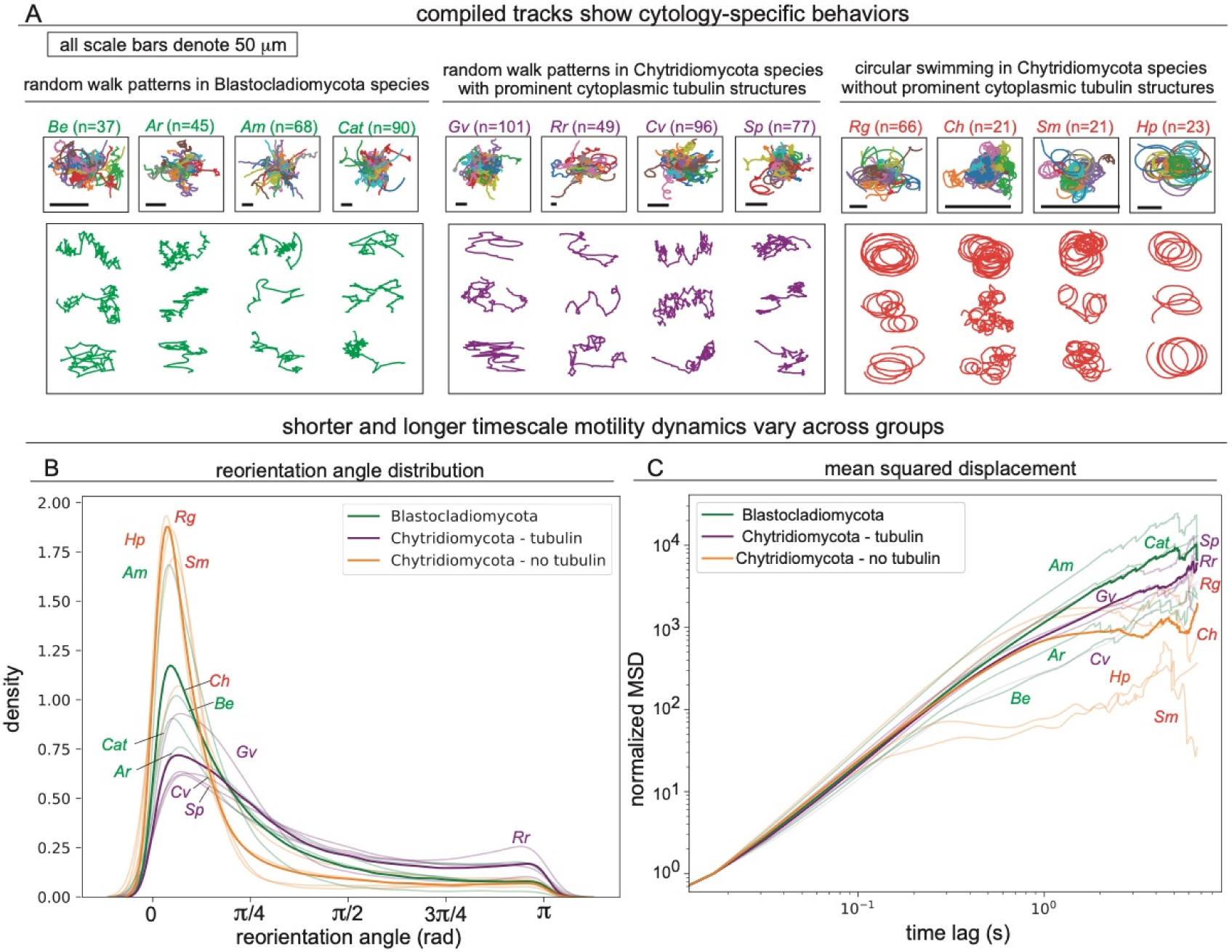
Movement tracks of zoospores of 12 zoosporic fungal species. (A) Movement tracks show cytology-specific swimming pattern behaviours, with all Blastocladiomycota species and Chytridiomycota species possessing prominent tubulin-based cytoplasmic structures swimming in a random walk pattern, while Chytridiomycota zoospores without prominent cytoplasmic tubulin structures swim in circular movements. Composite tracks are shown compiled into a single plot with starting points (time = 0) centred at the point of origin. (B) Distribution of turn (reorientation) angles for each group of zoospores (Blastocladiomycota species, Chytridiomycota species with prominent tubulin-based cytoplasmic components, and Chytridiomycota species with no prominent tubulin-based cytoplasmic structures). Species-averaged distributions are shown as partially transparent lines, as labelled. (C) Ensemble- and time-averaged mean-squared displacement (MSD) for each group of zoospores. Species-averaged MSDs are shown as partially transparent lines, as labelled. For effective comparison between groups, MSD plots are normalised by the initial value (at time lag 1) for each species. Unnormalized MSDs for each cell are shown in Figure S3B. Labels: *Catenophlyctis sp.: Cat, Blastocladiella emersonii: Be, Allomyces macrogynus: Am, Allomyces reticulatus: Ar, Synchytrium microbalum: Sm, Clydaea vesicula: Cv, Chytriomyces hyalinus: Ch, Rhizoclosmatium globosum: Rg, Homolaphlyctis polyrhiza: Hp, Rhizophlyctis rosea: Rr, Geranomyces variabilis: Gv,* and *Spizellomyces punctatus: Sp*.

### The relationship between cell structure and swimming pattern

To gain insights into the cellular mechanisms which putatively underpin these differences in swimming patterns, we analyse the intracellular organisation of zoospores. We performed fluorescent confocal microscopy with antibodies and cytological stains against α-tubulin, actin and lipid components of the zoospores from all 12 species studied (Figure 2). To further support our observations we reviewed previous electron microscope ultrastructural studies for each of these species^12–19^ (Table S1). These results suggest that observed variation in motility patterns segregate with cytological characters, specifically the presence of prominent cytoplasmatic tubulin structures, and not by higher taxonomic classifications (Figures 2, 3A).

**Figure 2.**
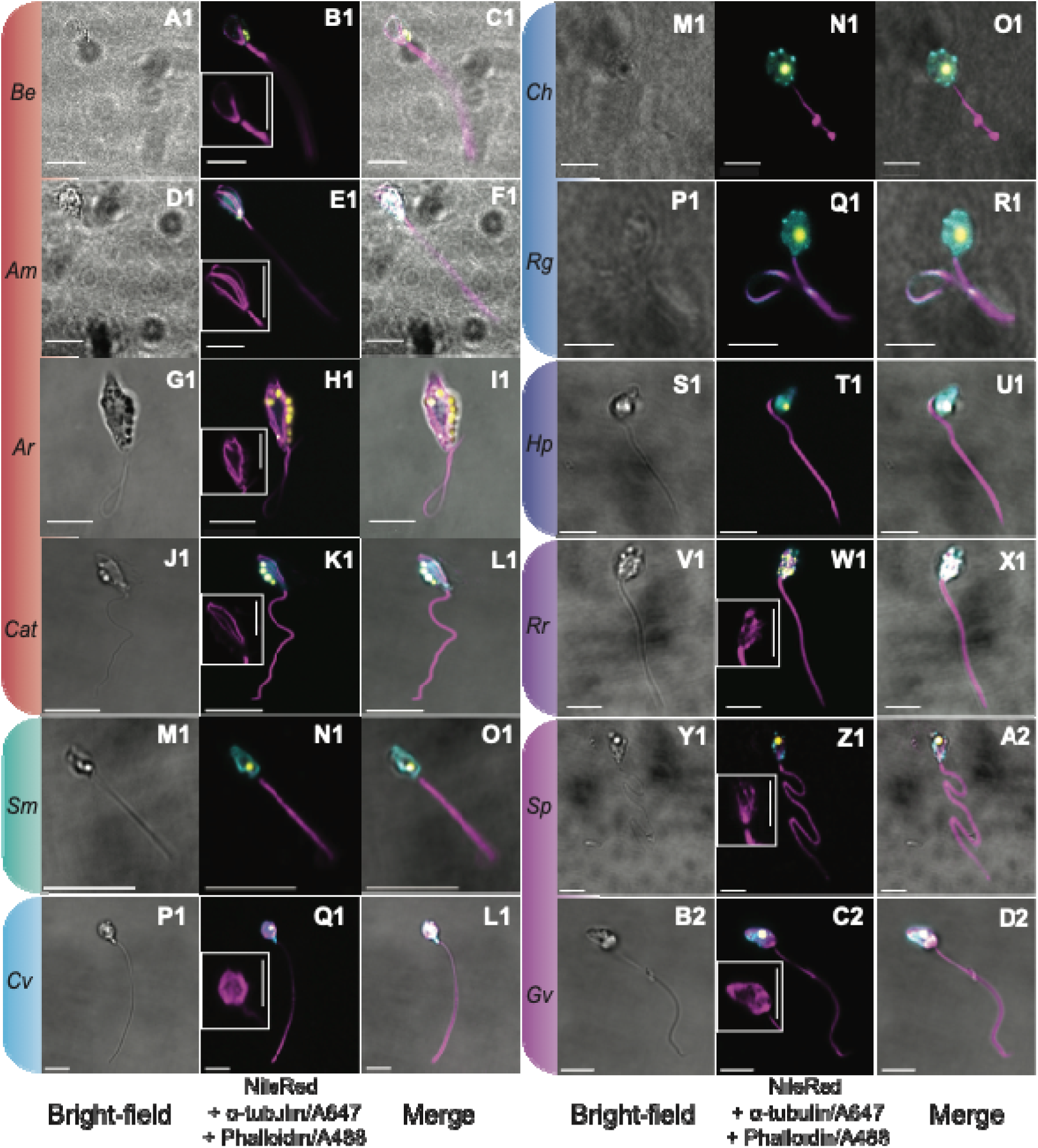
Ultrastructure of Blastocladiomycota and Chytridiomycota zoospores. First columns show brightfield phase contrast confocal microscopy images; second columns show fluorescent confocal microscopy micrograph of the zoospore; third columns show the merged images of brightfield and fluorescent channels. Microscopy images were taken and stained using the methods described in Galindo et al. (2022)^52^ with NileRed to stain lipid droplets, α-tubulin DM1A + Alexa Fluor 647 to stain tubulin and Alexa Fluor 488 Phalloidin to stain actin. In micrographs α-tubulin structures are stained magenta, actin stained cyan, and the lipid bodies (proposed to function as a light-sensing eye-spot) are stained yellow. In the fluorescent channel panels for those species which possess cytoplasmatic α-tubulin staining there is a small panel detailing the α-tubulin component. Species/micrographs bracketed/highlighted with red - belong to the Blastocladiomycota and those species micrographs with green-to-pink highlight-brackets correspond to Chytridiomycota from the: Synchytriales (M1-O1), Lobulomycetales (P1-L1), Chytridiales (M1-R1), Rhizophydiales (S1-U1), Rhizophlyctidiales (V1-X1) and Spizellomycetales (Y1-D2), respectively (the same colour code is used in Figure 3). Scale bars: A1-O1 = 10 µm; P1-D2 = 5 µm. Scale bars in small α-tubulin panels: B1, E1, H1 = 10 µm; K1, Q1, W1, Z1 and C2 = 5 µm. Labels: *Catenophlyctis sp.: Cat, Blastocladiella emersonii: Be, Allomyces macrogynus: Am, Allomyces reticulatus: Ar, Synchytrium microbalum: Sm, Clydaea vesicula: Cv, Chytriomyces hyalinus: Ch, Rhizoclosmatium globosum: Rg, Homolaphlyctis polyrhiza: Hp, Rhizophlyctis rosea: Rr, Geranomyces variabilis: Gv,* and *Spizellomyces punctatus: Sp*.

In all four Blastocladiomycota species, we observe random-walk movement patterns and show that the cytoplasmatic tubulin microtubules create prominent and complex cytoskeletal arrays or ‘ribs’ within the cell body which extend from the kinetosome of the flagellum around the periphery of the main body of the cell. Our microscopy shows that an actin cytoskeleton is much less apparent (Figure 2, 3A; n ≥ 10 cells observed for each species). Similarly, for two Chytridiomycota species that show random-walk movement patterns, *S. punctatus* and *G. variabilis* (Spizellomycetales) we also observe prominent tubulin microtubules that radiate at random into the zoospore cytoplasm from the kinetosome (n ≥ 10) i.e., forming rib-like structures similar to those seen in the Blastocladiomycota. For these Spizellomycetales, actin is present and forms patches in both species (the presence of these patches indicates possible actin organisation centres^20,21^).

Two additional Chytridiomycota species also showed prominent accumulations of non-assembled cytoplasmic α-tubulin-stained material; *R. rosea* and *C. vesicula*. Although these were not organised as clearly defined ribs in our micrograph analyses, such cell forms also performed random-walk movement behaviours. The zoospores of *R. rosea* (n = 10) showed both seemingly randomly arrayed α-tubulin accumulations and actin patches through-out the cytoplasm. Interestingly, microscopy imaging of *C. vesicula* identified an additional distinct α-tubulin organisation consisting of a peripheral cytoplasmic mesh of microtubule structures (n = 10). In *C. vesicula* we observe cytoplasmic actin co-localizing with the canonical vesicles found near the kinetosome of this species^15^ (Figure 2).

In contrast to the previously described species, we find no tubulin structures in the cytoplasm of *S. microbalum* (Synchytriales) and no major structures the other three Chytridiomycota: *H. polyrhiza* (Rhizophydiales), *C. hyalinus*, and *R. globosum* (Chytridiales) beyond a single microtubule structure connecting the base of the flagellum to the microbody-lipid globule complex organelle (MLC) only observed in TEM studies (n ≥ 10 cells observed for each species) (Figure 2, 3A, Table S1). Furthermore, in all four zoospores forms actin is the only detected cytoplasmic-cytoskeletal protein using our current confocal microscopy protocol. These four species swim in a circular pattern.

As for the lipid organelles, in Chytridiomycota we consistently observed one large lipid globule (n ≥ 10 – for each of these eight species) in their MLCs, with one exception found in the chytrid *R. rosea* in which we identified numerous small lipid droplets, a characteristic of this species^18^ (Figure 2). In the Blastocladiomycota we consistently observed clusters of several lipid globules (n ≥ 10 – for each of these four species), usually located towards the flagellum as part of the side-body complex organelle (SBC). In the case of *A. reticulatus* the lipid droplet clusters are numerous and run down the entire length of one side of the zoospore’s cell body (Figure 2).

Our video imaging results identified that the random walk patterns are specifically characterized by straight movements, followed by a stop and then a redirection of the cellular body. Observations of movement patterns for *B. emersonii* (*Be*) led to the hypothesis that re-direction occurs by a hinge-like bending of the area between the cytoplasmic microtubule and the flagellar kinetosome, and in an angle greater than the final angle of movement (Figure 3B). By performing live-fluorescent imaging of zoospores after staining with NileRed (lipid stain; n = 50) we observed that in *Be* the bend and subsequent redirection occurs to one side of the cell, the side where the SBC is located; suggesting this organelle is a constituent part and/or determinant of the hinge articulation which determines swimming redirection (Figure 3B).

**Figure 3.**
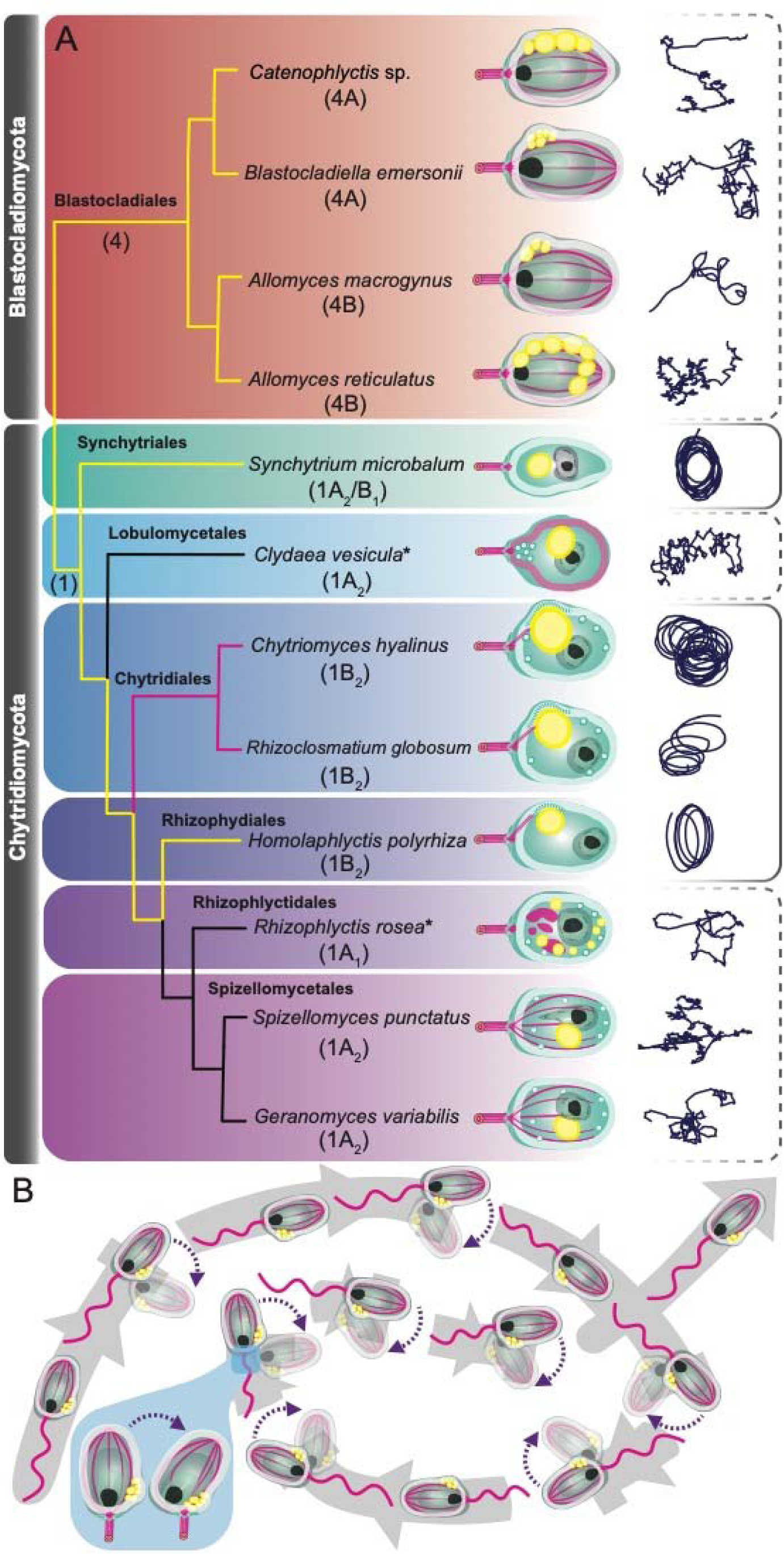
Depiction of motility, ultrastructure and phylogenetic relationships of the zoosporic fungal species from this study. (A) Phylogenetic relationships adapted from the multigene phylogeny from Amses et al. 2022^60^ showing the absence (black branches) or presence of homodimeric (yellow branches) or heterodimeric (magenta branches) RGCs opsin complexes adapted from Broser et al. 2023^59^. Drawings depict the overall ultrastructure of each zoospore including their lipid, actin, and α-tubulin-based components. A representative trajectory highlighting the overall movement pattern of each species is shown to the right of each drawing and grouped by a continuous (circular pattern) or discontinuous line (random-walk pattern). Number in parentheses bellow the species names and main branches indicate the lipid-organelle MLC classification of each zoospore according to Powell’s (1978) classification^45^. See Table S1 for further references. (B) Reconstruction based on our finding of the random-walk pattern of *B. emersonii* zoospores.

### Pharmacological inhibition demonstrates tubulin structures determines how random walker cells reorient

To test the involvement of tubulin cytoplasmatic structures in their swimming patterns, we treated representative species of each swimming behaviour and cytoplasmic cytoskeletal type with 1 µM of tubulin stabilizing (Taxol) and depolymerizing (Nocodazole) drugs. We selected *B. emersonii* (*Be*; Blastocladiomycota) and *S. punctatus* (*Sp*; Chytridiomycota) given their prominent microtubular ribs and marked random-walk movement, and *R. globosum* (*Rg*; Chytridiomycota) due to its lack of major α-tubulin cytoplasmic structures, circular swimming and its increasing utility as a model system^22^. We observed that when treating random-walk species *Be* and *Sp* with Nocodazole their overall swimming behaviour changes to manifest a circular swimming pattern (Figure 4A). However, there was no observable change in the swimming behaviour of the circular swimming *Rg* zoospores treated with nocodazole or taxol. Additionally, after imaging these nocodazole-treated and taxol-treated zoospores, we confirmed that nocodazole-treated zoospores lack their characteristic tubulin ‘rib’ cytoplasmatic arrangements, while there was no change in the tubulin arrangement of the taxol-treated species (Figure 4B-G). We also note that for the nocodazole-treated *Be* zoospores the lipid component (an organelle hypothesised to be key for the light sensing function^23,24^), which is usually located next to the base of the flagellum, loses its posterior cellular position and is relocated to random positions within the cell. Lastly, losing their tubulin cytoplasmic structures caused the zoospore cell morphology to change, from a tear-shape morphology towards a sphere-shape in *Be* and a reversed tear-shape in *Sp* (Figure 4C-F). Besides a slightly shorter flagellum in *Be*, we did not observe major changes in the tubulin structures of the flagellum across a range of drug treatments for the three fungal zoospores tested.

**Figure 4.**
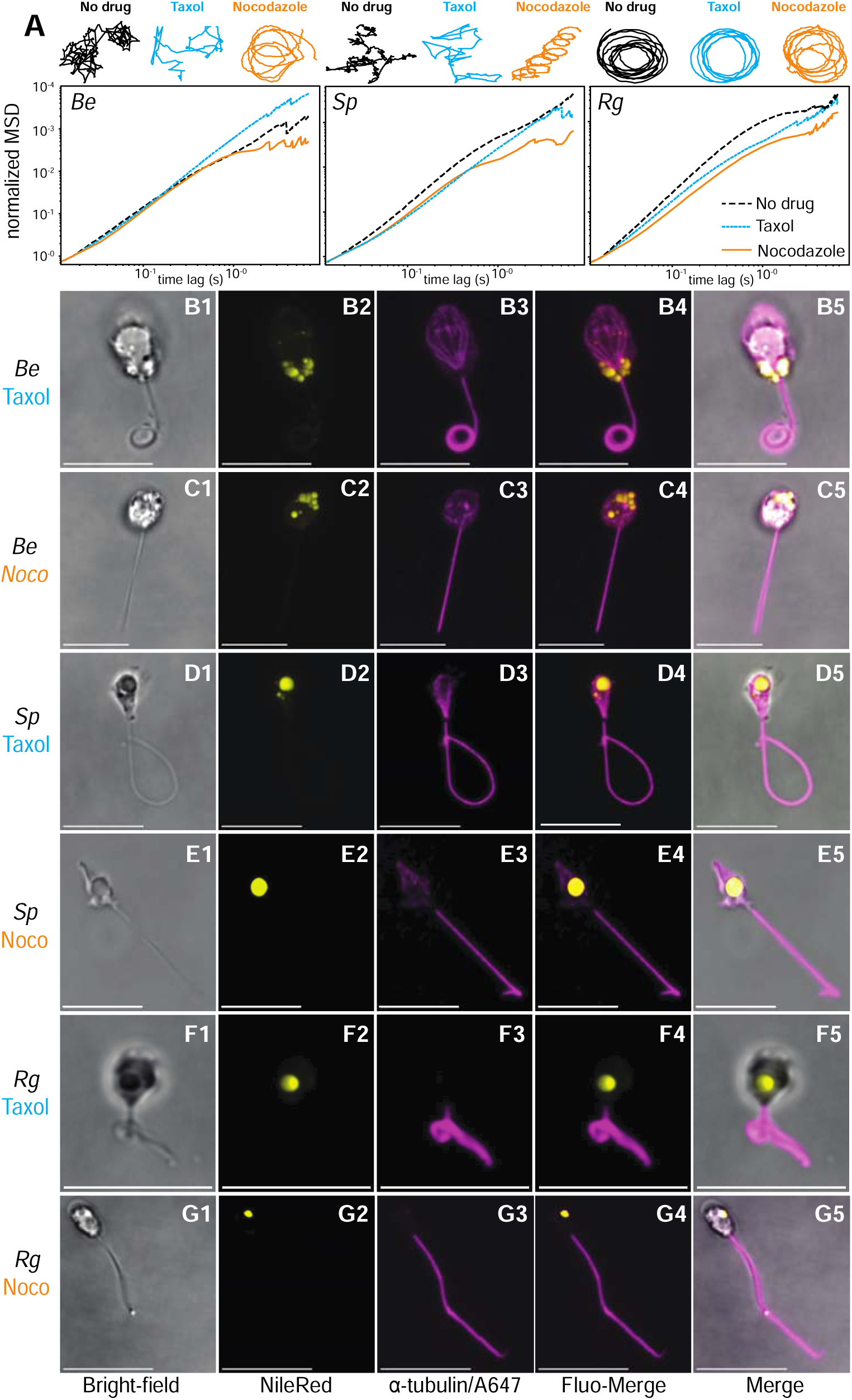
Movement tracks and confocal imaging of fungal zoospore from three species after treatment with Nocodazole and Taxol. (A) Representative swimming patterns and ensemble- and time-averaged mean-squared displacement (MSD) for each treatment in three zoospores fungal species. (B1-G5) Confocal imaging of drug treated zoospores. The first column shows brightfield phase contrast confocal microscopy images (B1-G1); second column shows in yellow the channel for NileRed staining of lipid droplets (B2-G2); third column shows in magenta α-tubulin DM1A + Alexa Fluor 647 staining of α-tubulin (B3-G3); fourth column shows the fluorescent channel merged (B4-G4); fifth column shows the merged images of brightfield and fluorescent channels (B5-G5). Scale bars = 10 µm. *Blastocladiella emersonii: Be, Spizellomyces punctatus: Sp, Rhizoclosmatium globosum: Rg*.

We performed an additional series of controls to test the involvement of actin in zoospore swimming behaviour by using actin depolymerizing drugs (Cytochalasin D, Latrunculin B and Jasplakinolide). As observed previously on *Batrachochytrium dendrobatidis*^25^ these drugs caused mass zoospore encystation in these three species, and no change in the swimming behaviour of the few remaining swimming zoospores was observed (Figures S4, S5).

## Discussion

### An association between zoospore swimming pattern and cytoplasmic tubulin structures

Swimming patterns have been studied in several unicellular eukaryotes (e.g.,^26–29)^. Zoospore swimming patterns in the protist *Phytophtora*, for example, have been shown to contribute to their dispersal, host colonisation and activity as plant pathogens^29,30^. These findings have led to the suggestion that chemical inhibition of motility could be used as a strategy for controlling the spread of pathogenic zoosporic species^29^. However, little is known about the swimming behaviour of fungal zoospores despite the crucial role of this life-cycle stage for dispersal and therefore host infection^5^.

Most chytrid species form uniflagellated zoospores, with only a few known multiflagellated species (e.g., *Neocallimastix* and *Piromyces*^31^) and lineages that form spores without flagella (due to a secondary loss in *Hyaloraphidium*^32^). Zoospores can move using two distinct mechanisms: swimming propelled by beating of the tubulin articulated flagellum^3,33^ or amoeboid crawling using actin driven pseudopodia^34,35^. These different locomotive modes are suited for specific behavioural goals; as such, there are fungal species which are both swimmers and crawlers (e.g., *Paraphysoderma*^35^), species with swimming zoospores which under certain conditions will switch to crawlers (e.g., *Batrachochytrium dendrobatidis*^20^) or with zoospores that exclusively crawl (e.g., Sanchytriomycota^34^). Broadly speaking, crawling is better adapted for traveling short distances on surfaces in liquid environments (e.g., exploration of trophic substrate); however, flagellum-based swimming is ideal for medium range dispersal in larger volumes of H_2_O^9^ (e.g., exploration of a water column to colonise a niche habitat). Thus, swimming plays an important role in facilitating the trophic life cycle of these fungi, a function that is often governed by chemo- or photo-taxis function^23,36–38^ to enable navigation towards, and colonisation of, niche habitats. These functions therefore drive the mycoloop^5,7,39^.

To our knowledge, this is the first systematic study of swimming patterns across a wide representation of the chytrid fungi with the patterns identified placed into a cellular and phylogenetic context (Figure 3A). Here we report two distinct swimming patterns among different fungal zoospore forming species: Chytridiomycota zoospores swim both in circular tracks (n = 4 spp), and in random-walk movement (n = 4 spp), while the Blastocladiomycota zoospores studied adopt a random-walk movement pattern (n = 4 spp). These patterns have previously been noted in a subset of chytrid fungi, providing additional data consistent with the pattern outlined here^40^. Specifically, circular swimming in the zoospores of the Rhizophydiales chytrid *Rhizophydium planktonicum*^41^ and random-walk swimming patterns in the zoospores of the Blastocladyomycota *Paraphysoderma sedebokerense*^35^ and *Physoderma* spp.^42^ from the order Physodermatales. These additional results suggest that a wider diversity of Blastocladiomycota species swim like the Blastocladiales studied here.

To address whether these swimming patterns form a consistent correlation with key cellular ultrastructure, we first perform fluorescence confocal microscopy of zoospores from all 12 species (Figure 2). Using this approach, we observed prominent cytoplasmic tubulin structures in all four Blastocladiomycota species and in four of the Chytridiomycota species analysed in this study which swim using the random walk patterns (Figure 3A). Similar to Blastocladiomycota, the Spizellomycetales Chytridiomycota fungi possess a complex tubulin-based cytoplasmatic structures with radiating microtubules (‘ribs’), as shown in previous ultrastructural TEM-based studies^16,19,43^ (Table S1). Despite also being prominent in our images, neither the cytoplasmatic unassembled tubulin structures observed here in the zoospores of *R. rosea,* nor the peripheral tubulin based cytoskeleton observed in *C. vesicula* were previously detected in TEM-based descriptions^15,18^. However, the strains of *R. rosea* (JEL0764) and *C. vesicula* (JEL0476) used, are different to the strains used for the original descriptions (Barr186/BR186 and JEL369 respectively^15,18,44^) so some variation in cytoskeletal structure may be present within these species complexes. *R. rosea* zoospores also have a large fibrillar rhizoplast extending from kinetosome which may fulfil a related function by acting as a hinge during the re-direction of movement^18^ (Table S1). Additionally, all these observations were made using TEM^15,18^, thus, these analyses would fail to detected non-electron dense structures, specifically failing to identifying dissociated tubulin material like the cytoplasmic tubulin structures detected in our analyses of these two species (Figure 2). The demonstration that some tubulin-based material can be observed in TEM, like the microtubules of Blastocladiomycota/Spizellomycetales (rib-like tubulin arrangements), while in others cytoplasmic tubulin can only be observed using fluorescent staining (*C. vesicula* and *R. rosea*); represents evidence of key differences between the cytoplasmic tubulin-based structures of these different zoosporic species. It is important to note that we refer to all these cytoplasmic tubulin structures as ‘prominent’ because we are aware that in the circular-swimming zoospores of *C. hyalinus*, *R. globosum* and *H. polyrhiza* there is a minor cytoplasmic tubulin component in the form of small bundles of microtubules connecting the kinetosome with the lipid-organelle’s rumposome (Figure 3A), which have also been observed using TEM^45^ (Table S1).

Taken together, our analyses confirm that the presence of prominent cytoplasmatic tubulin of various arrangements and densities correlates with the different swimming patterns observed. Specifically, zoospores from Chytridiomycota species that lack prominent cytoplasmatic tubulin structures swim using a circular pattern. In contrast, in all four Blastocladiomycota species and in those Chytridiomycota species in which there are prominent α-tubulin-stained structures within their cytoplasm, we observe a random-walk swimming pattern (Figure 3A).

Next, we sought to experimentally test the correlation between swimming behaviours and cytoplasmic tubulin structures by pharmacologically inhibiting or stabilizing intracellular tubulin using nocodazole and taxol, respectively. These drugs have successfully been used on several microbial eukaryotes to study cytoplasmic and flagellar tubulin-based structures^46–50^. Our results show that zoospores of fungal species with prominent cytoplasmic tubulin structures treated with nocodazole shifted their swimming patterns from random-walk to circular pattern (Figure 4). We note neither drug stopped flagellum swimming and in all three test species tubulin structures of the flagellum showed no major structural changes besides a putative and minor reduction in flagellar length in nocodazole-treated *Be* (Figure 4). Flagellar length does not seem to be factor determining the observed swimming patterns. All described zoospores vary in their flagellar length regardless of their swimming patterns, and species with the same flagellar length (e.g., ∼20 µm) can present either circular (e.g., *C. hyalinus*) or random-walk patterns (e.g., *S. punctatus*) (Table S1). In our test species *Rg* retains a long flagella and circular swimming across all treatments and after nocodazole treatment *Sp* retains its flagellar length, but the movement behaviour is transformed to a circular swimming mode. However, our results do demonstrate that we succeeded in inhibiting the polymerization/formation of ‘ribs’ within the main body of *Be* and *Sp* zoospores which became circular swimmers after treatment (Figure 4). Thus, we conclude that the capability of zoospores to perform ‘complex’ movements in a random walk fashion is linked to the presence of prominent cytoplasmatic tubulin structures. These findings demonstrate that a prominent tubulin-cytoplasmatic component may allow zoospores to swim using intricate patterns in which they can redirect movement by making sudden changes in direction; the random-walk pattern. These microtubular structures that radiate from the base of the flagellum may contribute to this pattern by forming a hinged articulation which bends the cell to re-orientate direction of travel towards a new direction of movement (Figure 3B). On the other hand, cells lacking tubulin cytoplasmic structures may restrict zoospores to swim using a constant circular/spiral-like pattern of movement.

### Is movement pattern connected to light-sensing and the lipid organelle in these zoospores?

Fungal zoospores possess different lipid organellar arrangement globally referred to as the microbody-lipid globule complex (MLCs) and are structures known to be involved in storage and production of chemical energy for flagellar beating^45^. There are different ‘types’ of MLCs which vary in structural arrangement across chytrid species. These structures are composed of microbodies, mitochondria, ribosomes and one or more large membrane-enclosed lipid globules (Figure 2, 3A)^2,45^. Using Powell’s (1978) classification^45^ all MLCs discussed here from Chytridiomycota are type 1, which can be classified as A or B; depending on the respective absence or presence of a rumposome (a cisterna-like disc of membranous tubules); zoospores can be further classified as A/B_1_ or A/B_2_ regarding the arrangement of different cellular components. The lipid-filled MLC organelles from Blastocladiomycota are type 4, and are alternatively classified as side-body complexes (SBCs) (Figure 2, 3A)^1,13,45,51^.

In addition to energetics, these organelles have been hypothesised to be involved in light perception^23,24,52,53^. Indeed, *Blastocladiella emersonii* and *Allomyces reticulatus* zoospores show positive phototaxis^23,37,54,55^. In particular, the SBC of *B. emersonii* was found to be in close spatial association with a layer of RGC (or CyclOp) proteins (a fusion protein composed of a type I microbial rhodopsin and guanylyl cyclase enzyme domain) which participates in a cGMP-mediated light-sensing pathway^23,56,57^ by controlling flagellar activation from its localization next to the SBC. Both these systems are in close physical association at the base of the flagellum. Recent analyses suggest that the molecular elements necessary for a functional RGC based light sensing pathway are present in the eight of the twelve fungal species analysed here^52,58,59^. These include six species with homodimeric RGC variants (all four Blastocladiomycota species, *S. microbalum* and *H. polyrhiza*) and two species with heterodimeric RGC complexes (Chytridiales: *C. hyalinus*, *R. globosum*) (Figure 3A), which consist of two subunits with distinct photochemical properties which evolved from an ancient duplication of the RGC protein^58,59^. We observe no association between RGC type, swimming pattern and ultrastructure. However, within the Chytridiomycota analysed here, those species that swim in circular patterns possess RGCs and MLCs type 1B and those with random-walk movement lack RGCs and have MLCs type 1A (Figure 3A), but wider sampling is needed to further test this pattern.

We see no direct correlation between number of lipid structures (one large lipid body vs multiple smaller lipid bodies) and either swimming pattern or possession of the heterodimeric RGC system. This result infers that the lipid body structure/morphology is not a factor in determining variant swimming behaviours or variance in light-perception mechanisms. However, some swimming behaviours may be influenced by the presence of the lipid structure. *Be* produces zoospores with the RGC system and that swim in a random-walk pattern and the SBC is found at one side of the kinetosome. Our experiments show that the cell-bending and redirection of motion in *Be* zoospores occurs always towards the side of the cell in which the SBC is present (Figure 3B). Among the possible explanations could be that prominent SBC is structurally restraining the movement of the *Be* zoospores towards that side, or that the zoospore moves towards the direction in which the light sensing organelle gets its stimulus. The position of MLCs relative to their movement is harder to assess with the random-walker Chytridiomycota species (e.g., *Sp*), since their MLCs are found in the centre of the cell and are unattached from the kinetosome. In circular swimming chytrids, small cellular sizes and large MLCs complicate the task of determining the position of MLCs relative to their circular motion, thus they remain to be characterized. Lastly, fluorescent confocal microscopy revealed that nocodazole-treated circular-swimming *Be* zoospores not only lacked cytoplasmatic tubulin microtubules, but they also loss the typical arrangement of the SBC organelle at the base of the flagellum, providing further evidence that both structures interact through the rib-like tubulin structures (Figure 4C). The physical link between the SBC and the base of the kinetosome may be essential for the transduction of the signal to flagellar movement which then translates to the observed swimming pattern.

Regarding a light-perception/swimming pattern relationship, we show that *A. macrogynus* (*Am*) tends to swim in more regular, directed trajectories than other random-walkers; as shown by its narrower turn angle distribution and its MSD scaling with higher exponent (Figure 1B-C). One possible explanation is that *Am* is the only Blastocladiomycota in our sample known to have replaced light sensing with chemo sensing^37^. Thus, if our set up is triggering light sensing (all observations are under white light) but not chemo sensing (no amino acid gradients), we could hypothesize that light sensation is constantly triggering reorientation in random walker Blastocladiomycota except in *Am*, which has no response to light. Overall, these experiments point towards a link between zoospore ultrastructure, movement, and the light sensing organelle component. Our findings provide comparative model systems for understanding the link between cytology, sensing and divergent motility behaviour in single-celled eukaryotic microbes.

## Acknowledgements

This work was funded by the Horizon 2020 research and innovation programme under the European Marie Skłodowska-Curie Individual Fellowship H2020-MSCA-IF-2020 (grant agreement no. 101022101—FungEye) and Leverhulme Trust Project Grant (RPG-2022-234; The evolutionary diversification of a sub-cellular fungal eye). T.A.R. is supported by a Royal Society URF (URF/R/191005). We gratefully acknowledge the Micron Advanced Bioimaging Facility (supported by Wellcome Strategic Awards 091911/B/10/Z and 107457/Z/15/Z). We also acknowledge Suely Lopes Gomes (Universidade de São Paulo) for inspiring work on chytrid light perception. J.A.N. acknowledges support from All Souls College, Oxford and the National Science Foundation through the Center for Living Systems at the University of Chicago (grant no. 2317138). We are also grateful for the comments of three anonymous reviewers, which have improved this manuscript from its original form.

## Author contributions

L.J.G., T.A.R. and J.N.: conceived the study; L.J.G.: microscopy imaging, drug assays and manuscript writing; T.A.R. manuscript writing; J.N.: imaging analyses and manuscript writing. All authors gave final approval for publication and agreed to be held accountable for the work performed therein.

## Declaration of Interests

These authors declare no competing interests.

## STAR Methods

### Resource availability

#### Lead contact

Further information and requests for resources and reagents should be directed to and will be fulfilled by the Lead Contact, Jasmine A. Nirody.

## Materials availability

This study did not generate any new or unique reagents.

## Data and code availability

All data are available in the figures, tables, and data files associated with this manuscript. Plots of all tracks for each species are shown in Supplementary Material. Raw videos are available at https://figshare.com/account/home#/projects/207943 and tracking data, detailed protocols, and analysis codes are provided at http://github.com/jnirody/fungalzoospores. Raw confocal microscopic images can be found at: https://figshare.com/projects/FungEye/132734. Any additional information required to reanalyse the data reported in this work paper is available from the Lead Contact upon request.

## Experimental model and subject details

Cultures of *Blastocladiella emersonii* ATCC 22665 (ATCC; American Type Culture Collection), *Catenophlyctis* sp. JEL0575 (CZEUM; Collection of Zoosporic Eufungi at University of Michigan; 18S rDNA sequence could indicate it belongs to a species of *Catenaria*^44^), *Clydaea vesicula* JEL0476 (CZEUM), *Chytriomyces hyalinus* CBS 675.73 (previously referred as *C. confervae* given that CBS 675.73 = ATCC 24931 = Barr 97 and Barr 97 is *C. hyalinus* ^44,61^; Westerdijk Fungal Biodiversity Institute), *Homolaphlyctis polyrhiza* JEL0142 (CZEUM), *Rhizophlyctis rosea* JEL0764 (CZEUM), *Geranomyces variabilis* JEL0559 (CZEUM), and *Spizellomyces punctatus* SW1 (CZEUM) were vegetatively grown in 25 cm2 culture flasks (Sarstedt) filled with 25 mL of PYG liquid media (0.13% w/v peptone, 0.13% w/v yeast extract, 0.3% w/v glucose). *Rhizoclosmatium globosum* JEL0800 (CZEUM) vegetative cells were grown on PYG agar plates (0.13% w/v peptone, 0.13% w/v yeast extract, 0.3% w/v glucose, and 1.5% w/v agar). *Allomyces macrogynus* Australia_3 (CZEUM) and *Allomyces reticulatus* ATCC 42465 (ATCC) was grown in 25 cm^2^ culture flasks filled with 25 mL of Emerson YpSs/4 liquid media (0.1% w/v yeast extract, 0.375% w/v soluble starch, 0.025% w/v dipotassium phosphate, 0.01% w/v magnesium sulfate). *Synchytrium microbalum* JEL0517 (CZEUM) vegetative cells were grown on PYG agar plates (0.13% w/v peptone, 0.13% w/v yeast extract, 0.3% w/v glucose, and 1.5% w/v agar) and in PYG liquid media (0.13% w/v peptone, 0.13% w/v yeast extract, 0.3% w/v glucose). All fungal vegetative growth was performed at 21°C and transferred every two weeks by inoculating 25 µl of previous culture to a new flask/plate containing 25 ml of media.

To induce sporulation of *B. emersonii* ATCC 22665, *C. vesicula* JEL0476, *C. hyalinus* CBS 675.73, *H. polyrhiza* JEL0142, *R. rosea* JEL0764, *G. variabilis* JEL0559, and *S. punctatus* SW1, 200 mL of liquid PYG were inoculated with 20 ml of vegetative cells in 175 cm^2^ culture flasks (Sarstedt), and incubated, with 150 rpm agitation, for 24 h at 21°C. To induce sporulation in *A. macrogynus* Australia_3, *A. reticulatus* ATCC 42465 and *Catenophlyctis* sp. JEL0575 vegetative growth was performed for ∼5 days in 25 ml of liquid YpSs/4 (*Allomyces*) or PYG (*Catenophlyctis*). Sporangia were then separated from the culture using sterile tweezers, washed with distilled H_2_O, and placed in 20 mL of distilled H_2_O in a 25cm^2^ culture flask at 21°C overnight. *R. globosum* JEL0800 and *S. microbalum* JEL0517 zoospore obtention was performed by growing this strain on PYG agar plates for 4 days. Plates were then flooded with 7 mL of distilled H_2_O and after resting for 15 min to allow sporulation, zoospore-containing H_2_O was transferred to a 15 ml falcon tube. Zoospores were then separated from sporangia using a 20 µm pluriStrainer (pluriSelect) into 50 ml falcon tubes. All sporulation inductions were performed at 21°C.

## Method details

### Images acquisition

Out of the 12 obtained liquid zoospore solutions, seven were PYG-based and five were H_2_O -based (*A. macrogynus*, *A. reticulatus*, *Catenophlyctis* sp., *S. microbalum* and *R. globosum*). After sporulation, both 7 μl and 100 μl of zoospore suspension aliquots were placed respectively on a slide and coverslip or placed on the bottom of the well of a flat bottom 86- well plate. *S. microbalum* was recorded only on plates and *A. reticulatus* culture died before plate videos could be recorded. Thus, we image zoospores both in a physically constrained and non-constrained environment; we observed no difference between the two conditions (Figure S6A). Alternatively, Nile-red (Thermo Fisher Scientific, Cat# N1142) was added in a 1:500 v/v concentration for 5 min before live-imaging of zoospores. Cells were imaged on an Olympus CKX53 inverted microscope with 10X, 20X, and 40X objectives according to the size of the zoospores and/or their swimming trajectories. Videos of 15 to 30 seconds were taken at 60 fps with a Std Chromyx HD camera mounted on a U-TV0.5XC-3-8 0.5x C-Mount Adapter. Exposures, focus, and stage position were kept constant while recording for motility analyses. The number of trials *n* recorded for each species is shown in Figure 1A.

Both PYG media and H_2_O have similar physical conditions and the use of different liquid media did not seem to affect the swimming behaviours of closely related species with similar cellular structures (e.g., circular swimming: *R. globosum* in H_2_O and *C. hyalinus* in PYG medium; random walk swimming: *A. reticulatus* in H_2_O and *B. emersonii* in PYG medium). We note that given the small size of the zoospores, minor changes in media viscosity are unlikely to affect cell mechanics. However, it is possible that environmental chemistry and the presence of different ion concentrations in the aqueous environment may modify swimming behaviour^62,63^. To explore this source of variation further, we re-recorded the swimming behaviour of zoospores of the 7 species which were originally recorded in PYG media also in distilled H_2_O media (by filtering and re-suspension in dH_2_O). We observed no change in the swimming patterns when moved to alternative media (Figure S6B).

### Tracking and motility analysis

Movies of swimming zoospores were viewed in FIJI^64^, and movement was tracked using a semi-automated protocol with the TrackMate plugin^65^. Before tracking, videos were pre- processed as follows: (1) the background was subtracted; (2) artifacts were detected and removed by subtracting the median image of the video from each frame. Cells were segmented using TrackMate’s Difference of Gaussian detector; manual quality control by eye was performed on the first frame for each video. The centre-of-mass of segmented images were joined to create tracks using a Linear Assignment Problem (LAP) tracker; maximum gap closing was set at 50 pixels and maximum frame gap at 10 frames. Final quality control of all tracks was performed using TrackScheme to ensure each track consisted of only one detected spot per frame.

### Drug assays

Drugs assays were conducted by exposing cultures of *B. emersonii*, *S. punctatus* (grown overnight) and *R. globosum* (grown for 4-days) for 5 hours at 21°C to 1 µM of Nocodazole (tubulin polymerization inhibition) and separately Taxol (microtubule stabilizing) solution resuspended in DMSO. Zoospores were strained-filtered from sporangia as described before and then fixed in 4% *w/v* paraformaldehyde (for confocal imaging) and/or video recording. Aliquots of zoospore solution of these three species were also exposed for 10 min at room temperature to 1 μM Latrunculin B (a drug which sequesters actin monomers), 100 μM Cytochalasin D (a drug which caps actin filaments), 10 μM Jasplakinolide (a drug which stabilizes actin filaments) solutions and then resuspended on DMSO for movement behaviour video recording. Protocols and drug concentrations were based on previous studies using these same drug treatments on either zoosporic fungi and/or alternative microbial eukaryotes^25,46–50,66^.

### Quantification and statistical analysis

Plotting of tracks and quantitative motility metrics were performed from exported COM data using in-house scripts written in Python. Random walk and circular motility patterns were quantitatively differentiated using reorientation angle (or turn angle), a measure of local cellular behaviour, and mean square displacement (MSD), a measure of the deviation of the position of a cell *r* over time. Turn angles at time *t* were calculated as: δΘ = tan - 1 [v(t) × v(t + 1)) / (v(t)• v(t + 1)]. The MSD of a cell at time *t* is calculated by MSD(r) = < r(t + r) - r(t) >, as a function of lag time r. MSD values are calculated across the entire trajectory for lag times τ ranging from 1 to 800 frames (0.017 to 13.3 s). Cells exhibiting circular motility patterns are expected to show plateaus at long-time MSDs. Plots of time-averaged MSD for each cell are shown as grey lines in Figure S16; ensemble-averages are shown for each species in red. For interspecies comparisons (Figure 1C), ensemble-averaged MSDs for each species were normalised by dividing by the MSD value at r *= 1*.

### Fluorescence assays

In all cases, zoospores were collected from the suspension medium by centrifugation at 1000 x g for 5 min followed by removal of the supernatant in order to concentrate the cells. In all cases the cell pellet was fixed in 0.5 ml of 4% *w/v* paraformaldehyde in 1X PBS and transferred into a 15 ml Falcon tube for 15 min at room temperature. Cells were concentrated by centrifugation and resuspended in PBS for the first washing step. The cell pellet was then resuspended and permeabilized in 0.5 ml of PBS containing 0.1% *v/v* Triton X-100 in PBS. A second washing step in 0.5 ml of PBS was performed and then the cells were blocked with 0.5 ml of 1% *w/v* BSA in PBS and incubated for 45 min at room temperature, followed by the addition of the primary antibody α-tubulin DM1A (Sigma-Aldrich, Cat# T6199, RRID: AB_477583) at a concentration of 1:500 v/v in 1% *w/v* BSA in PBS for 180 min at room temperature. After a third washing step of the primary antibody with 1X PBS, the secondary antibody Alexa Fluor 647 Goat anti-mouse IgG1 antibody was added (Thermo Fisher Scientific, Cat#A-21240, RRID: AB_2535809) to the PBS-resuspended fixed zoospore solution for 60 min at room temperature. Alexa Fluor 488 Phalloidin (Invitrogen, Cat# A12379, RRID: AB_2759222) and Nile Red (Thermo Fisher Scientific, Cat# N1142) were also added at a 1:500 v/v concentration for 60 min at room temperature. After two washing steps with 1X PBS, the final cell pellets were resuspended in 100 µl of 20% v/v Citifluor AF2 Antifadent Mountant Solution and 7 µl was placed on a slide and covered with a coverslip, which was then sealed with transparent nail polish on the edges to avoid evaporation. Cells were imaged on a Zeiss LSM-780 inverted high-resolution laser scanning confocal microscope with a Ph3 ×100 oil objective. Exposures were kept constant during experiments, and images collected using ZEISS ZEN Software (ZEN Digital Imaging for Light Microscopy), and analysed/formatted with Fiji ^64^.

## Notes

https://doi.org/10.6084/m9.figshare.21866169.v1

http://github.com/jnirody/fungalzoospores

### Competing Interest Statement

The authors have declared no competing interest.

### Summary of Updates

We have added more species across phyla to put our results in a stronger phylogenetic context (Figure 3) and performed some experiments with actin and tubulin depolymerizing drugs to test our hypothesis about the relationship between cytology and motility patterns (Figure 4).

