## Supplementary Figures and Tables for "Evolutionarily diverse fungal zoospores show contrasting swimming patterns specific to ultrastructure"

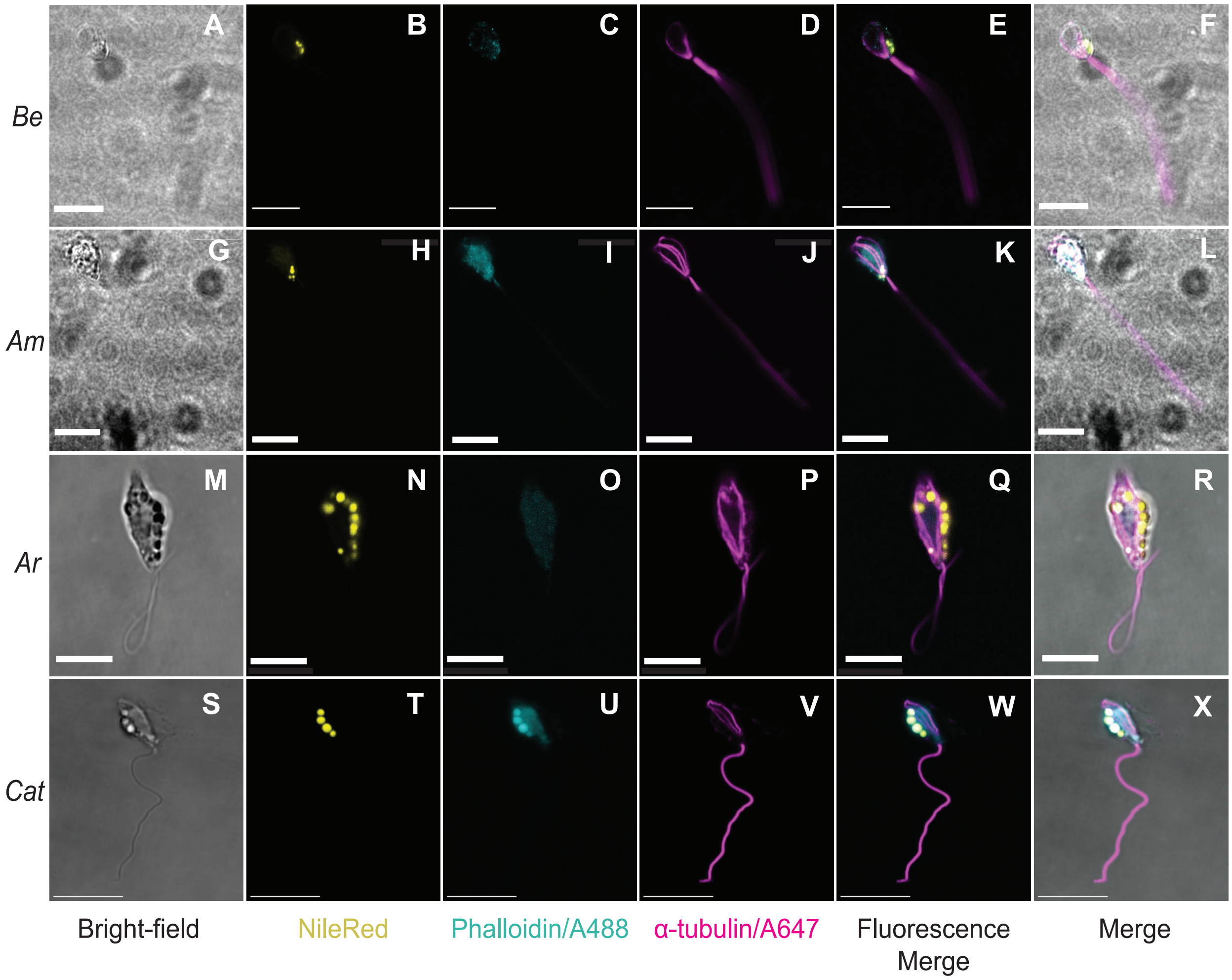


**Figure S1. Imaging of ultrastructure in Blastocladiomycota zoospores.** First columns show brightfield phase contrast confocal microscopy images (A, G, M, S). Second columns show fluorescent confocal microscopy micrograph of the zoospore stained with with NileRed to stain lipid droplets (B, H, N, T; yellow). Third column shows zoospores stained with Alexa Fluor 488 Phalloidin to stain actin (C, I, O, U; cyan). Fourth column shows α-tubulin DM1A + Alexa Fluor 647 staining for α-tubulin (D, J, P, V, magenta). The fifth column shows the merge of all fluorescent channels (E, D, J, P, V). The sixth column shows the merged images of brightfield and fluorescent channels (F, L, R, X). Scale bars: A1-O1 = 10 μm; P1-D2 = 5 μm. Scale bars in small α-tubulin panels: B1, E1, H1 = 10 μm; K1, Q1, W1, Z1 and C2 = 5 μm. Labels: *Catenophlyctis* sp.: *Cat, Blastocladiella emersonii*: *Be, Allomyces macrogynus*: *Am, Allomyces reticulatus: Ar*.

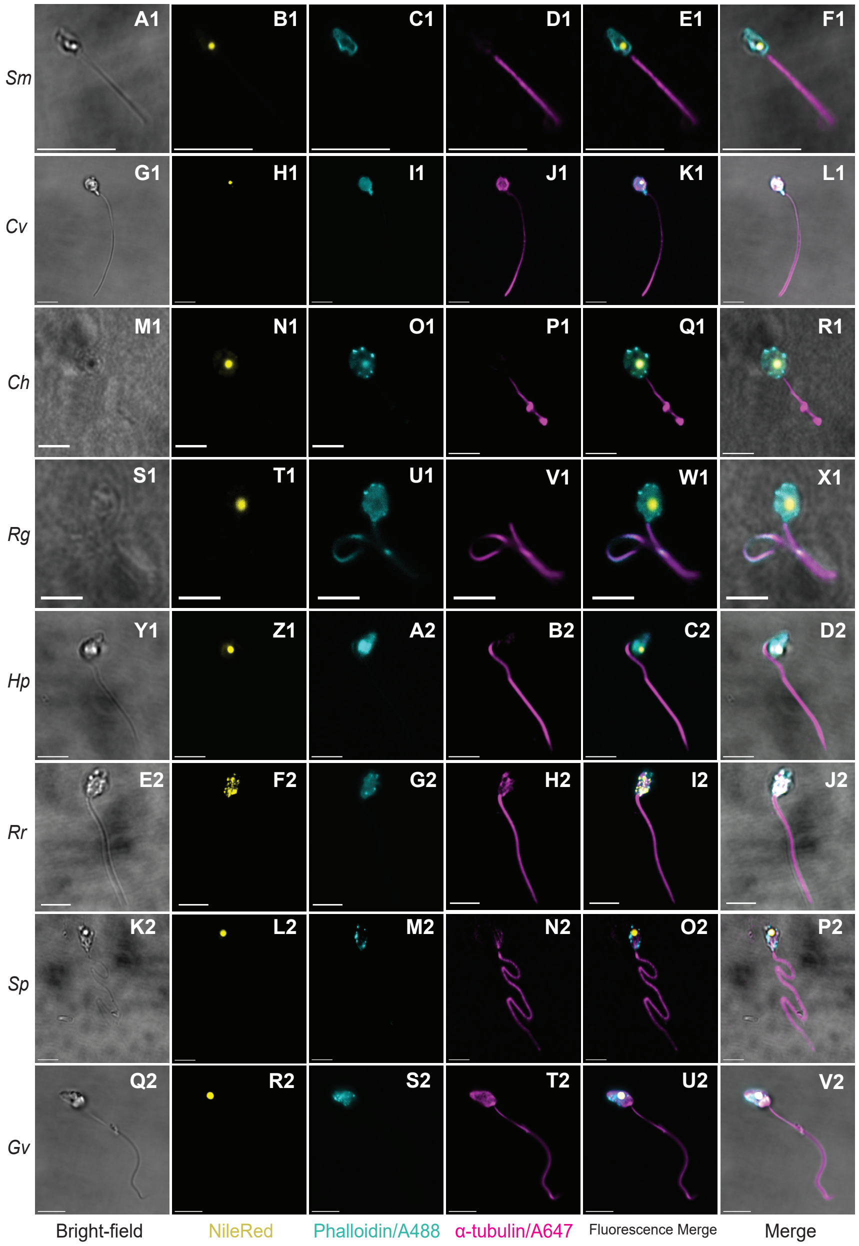


**Figure S2. Imaging of ultrastructure in Chytridiomycota zoospores.** First columns show brightfield phase contrast confocal microscopy images (A1, G1, M1, S1, Y1, E2, K2, Q2). Second columns show fluo- rescent confocal microscopy micrograph of the zoospore stained with with NileRed to stain lipid droplets (B1, H1, N1, T1, Z1, F2, L2, R2; yellow). Third column shows zoospores stained with Alexa Fluor 488 Phalloidin to stain actin (C1, I1, O1, U1, A2, G2, M2, S2; cyan). Fourth column shows α-tubulin DM1A + Alexa Fluor 647 staining for α-tubulin (D1, J1, P1, V1, B2, H2, N2, T2; magenta). The fifth column shows the merge of all fluo- rescent channels (E1, K1, Q1, W1, C2, I2, O2, U2). The sixth column shows the merged images of brightfield and fluorescent channels (F1, L1, R1, X1, D2, J2, P2, V2). Scale bars: A1-O1 = 10 µm; P1-D2 = 5 µm. Scale bars in small α-tubulin panels: B1, E1, H1 = 10 µm; K1, Q1, W1, Z1 and C2 = 5 µm. *Synchytrium microbalum: Sm, Clydaea vesicula: Cv, Chytriomyces hyalinus: Ch, Rhizoclosmatium globosum: Rg, Homolaphlyctis polyrhiza: Hp, Rhizophlyctis rosea: Rr, Geranomyces variabilis: Gv, and Spizellomyces punctatus: Sp.*

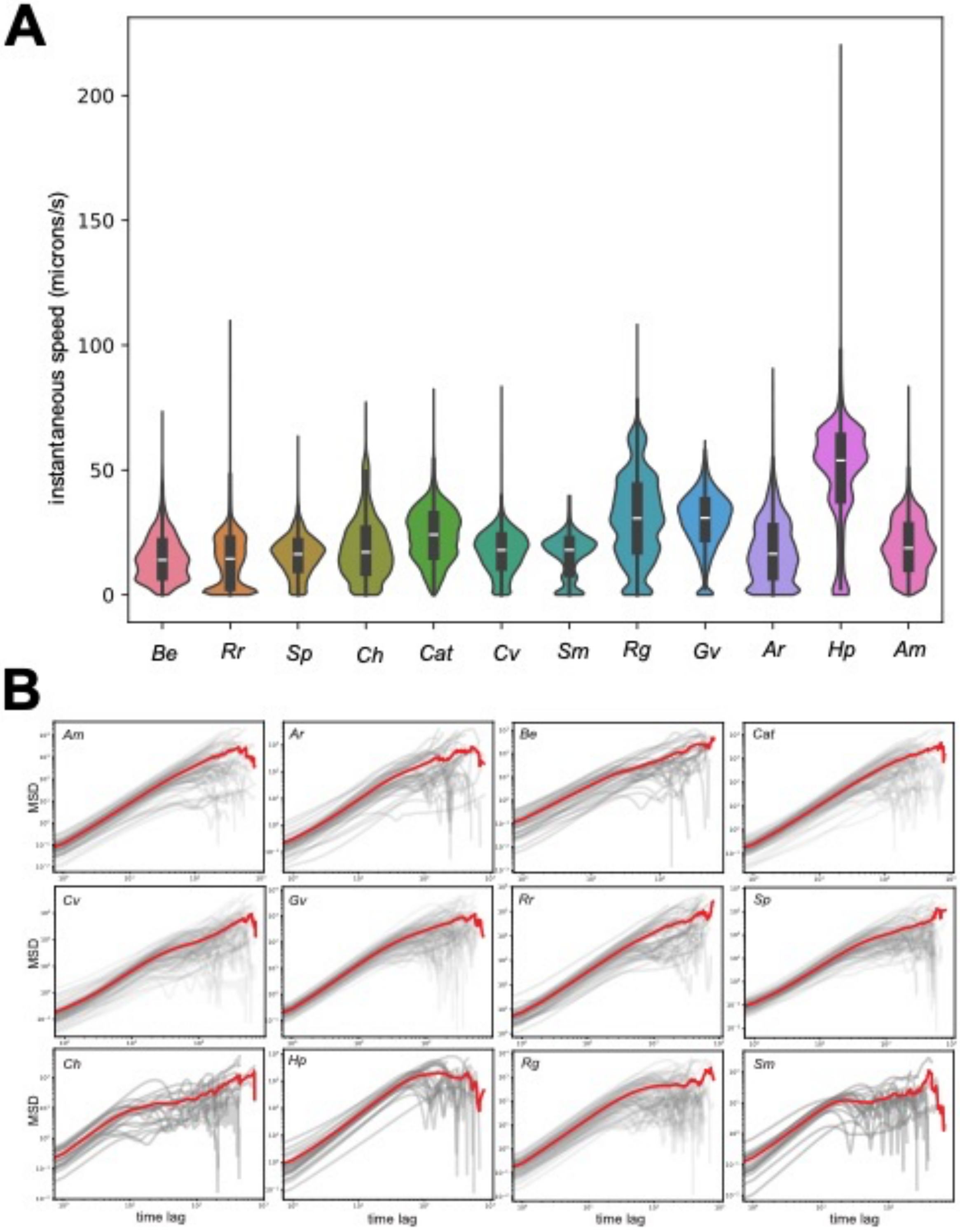


**Figure S3. (A)** Instantaneous speeds (in units of microns per second) over all trials for each fungal zoospore species. **(B)** Time averaged MSD values (grey lines) and ensemble averaged MSD values (red lines) for each fungal zoospore species.

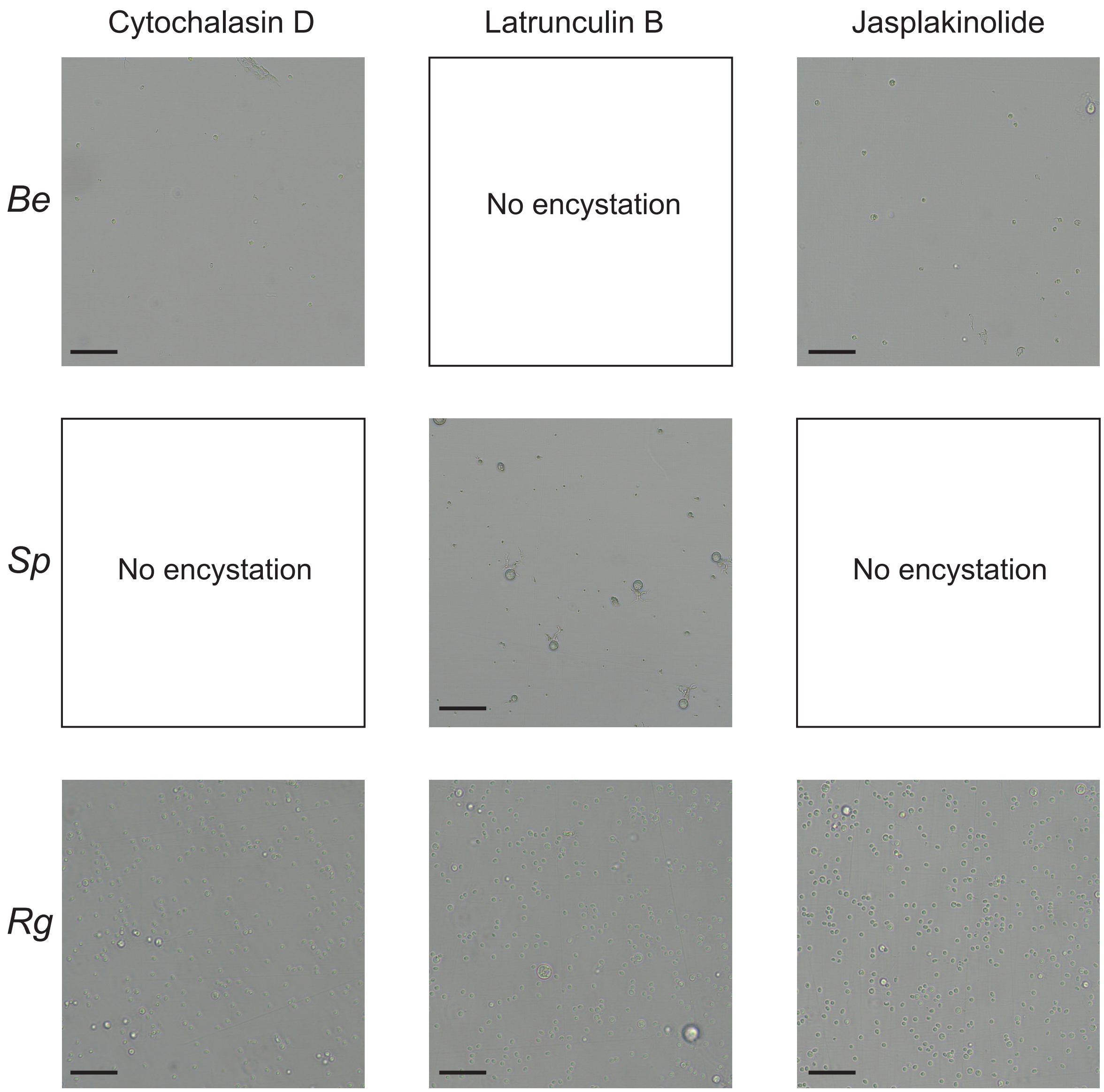


**Figure S4**. Encystation in zoospores from three fungal species after exposure to actin depolymerizing drugs. Zoospore solution of these three species was alternatively exposed for 10 min at RT to 1 μM Latrunculin B (to sequester actin monomers), 100 μM Cytochalasin D (to cap actin filaments), 10 μM Jasplakinolide (to stabilize actin filaments) solutions resuspended on DMSO before being recorded. Scale bars = 50 μm*. Blastocladiella emersonii: Be, Spizellomyces punctatus: Sp, Rhizoclosmatium globosum: Rg*.

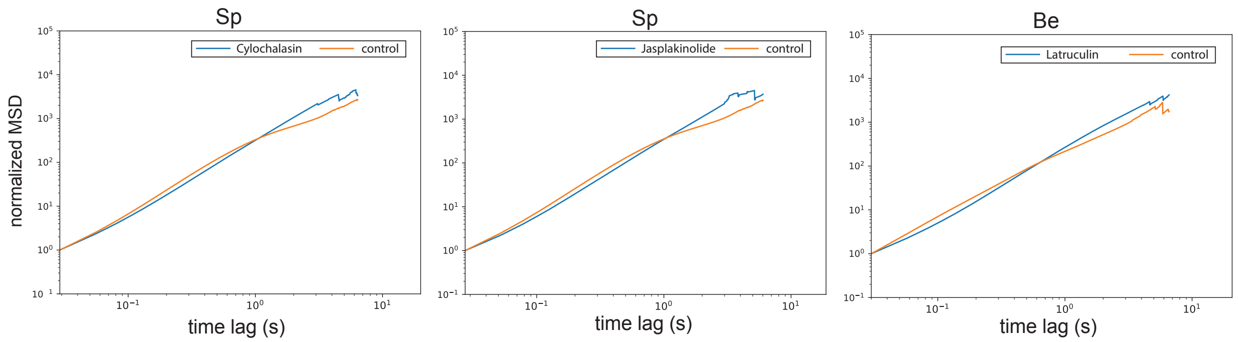

**Figure S5.** Time- and ensemble-averaged MSDs comparing motility patterns after exposure to actin depolymerizing drugs. *Blastocladiella emersonii: Be, Spizellomyces punctatus: Sp.*

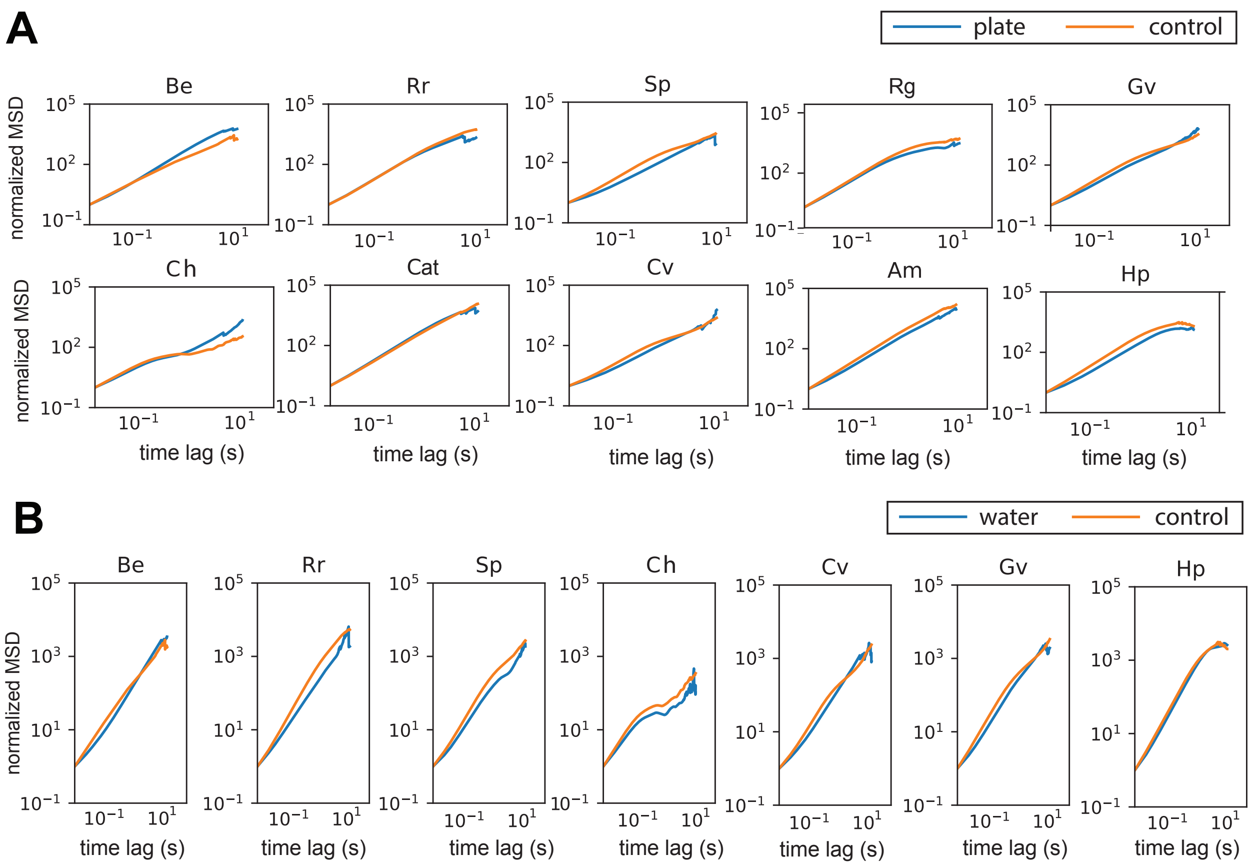


**Figure S6.** Time- and ensemble-averaged MSDs comparing the motility dynamics of various zoospore species in (A) an unconfined environment (86-well plate) and (B) water as opposed to a control condition in between two coverslips in media as denoted in the main text.

| **Species** | Swimming pattern | Cell size (wide-long; µm) | Cell shape | Flagellum length (µm) | MLC type (Powell, 1978) [1] | Rumposome | Cytoplasmic microtubule | Flagellar-MLC microtubular bundle | Reference cell diagram/TEM micrograph |
|---|---|---|---|---|---|---|---|---|---|
| *Catenophlyctis* sp. | Random-walk | 2-4.5[2] | Ovoid[2] | 25-39[2] | 4A (SBC)[1,3] | No[3] | Yes[3] | No[3] | [4] 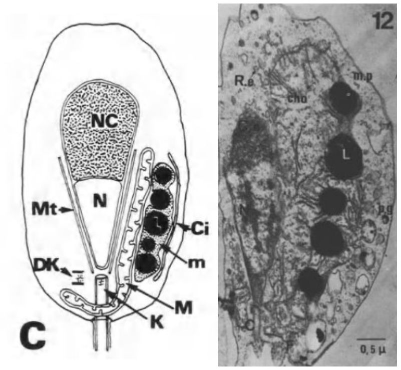  |
| *Blastocladiella emersonii* | Random-walk | 7-9[5] | Ovoid[5] | ~15[6] | 4A (SBC)[1,7] | No[5,7] | Yes[5,7] | No[5,7] | [5,8] 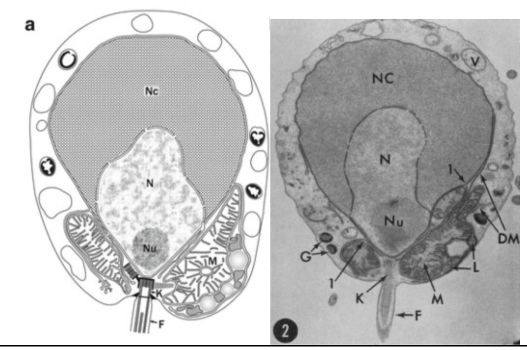  |
| *Allomyces macrogynus* | Random-walk | ~8-10[9] | Ovoid[10] | ~30[11,12] | 4B (SBC)[1,9,10,12] | No [9,10,12] | Yes [9,10,12] | No [9,10,12] | [12,13] 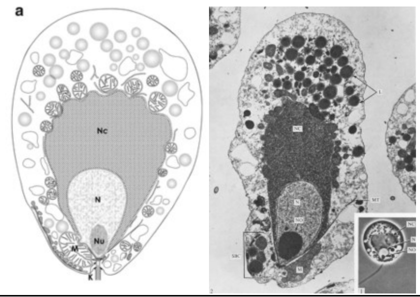  |

| | | | | | | | | | |
|---|---|---|---|---|---|---|---|---|---|
| *Allomyces reticulatus* | Random-walk | 8-12[14] | Ovoid[14] | ~31-35[14] | 4B (SBC)[1,14] | No[14] | Yes[14] | No[14] | [14] 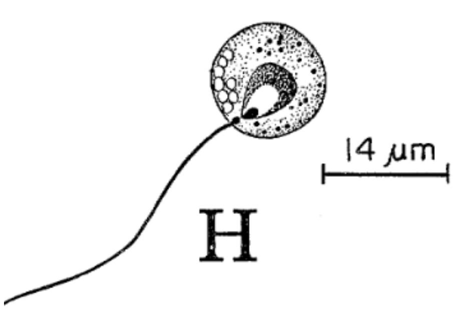  |
| *Synchytrium microbalum* | Circular | 2-3.5[15] | Elongated[15] | ~10[15] | 1$A_2$ or 1$B_1$[1,15] ($B_1$ since some species have a rumposome) | No[15] | No[15] | No[15] | [15] 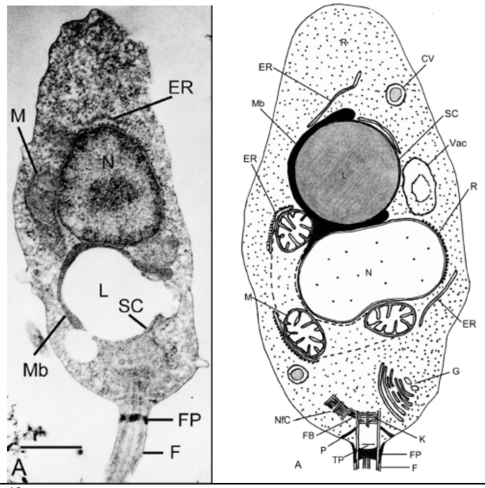  |
| *Clydaea vesicula* | Random-walk | ~5[16] | Spherical[16] | 20[16] | 1$A_2$[16] | No[16] | No[16] | No[16] | [16] 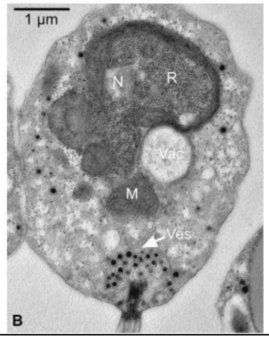  |

| | | | | | | | | | |
|---|---|---|---|---|---|---|---|---|---|
| *Chytriomyces hyalinus* | Circular | 3.5-6 [17,18] | Subspherical [17,18] | 18-20[17,18] | $1B_2$[1,17,18] | Yes[17,18] | No[17,18] | Yes[17,18] | [18,19] 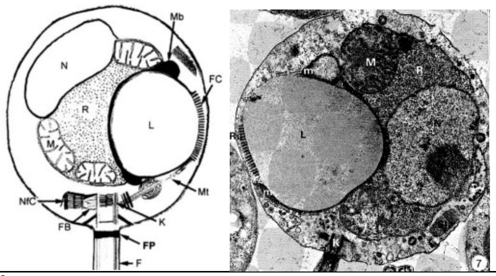  |
| *Rhizoclosmatium globosum* | Circular | 3-6[18] | Ovoid[18] | ~23[18,20] | $1B_2$[1,18] | Yes[18] | No[18] | Yes[18] | [18] 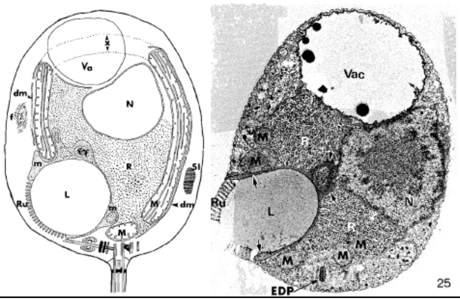  |
| *Homolaphlycti s polyrhiza* | Circular | 3.5-4.5[21] | Spherical[21] | ~28[21] | $1B_2$[21] | Yes[21] | No[21] | Yes[21] | [21] 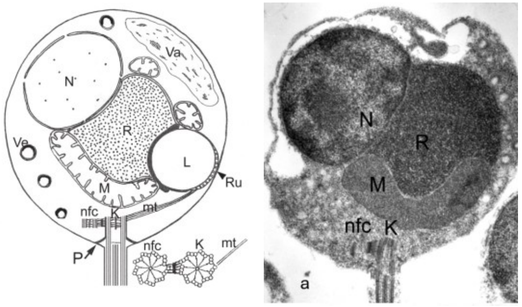  |
| *Rhizophlyctis rosea* | Random-walk | 4-5[22,23] | Spherical[22,23] | 20[22,23] | $1A_1$ [1,22,23] | No[22,23] | No, but large fibrillar rhizoplast[22,23] | No[22,23] | [22] 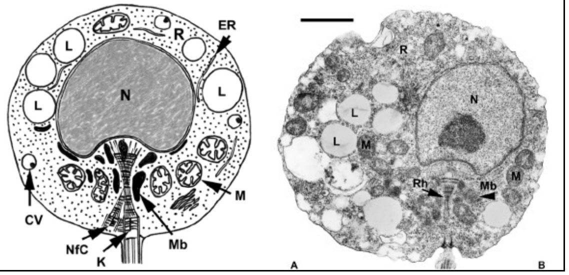  |

| | | | | | | | | | |
|---|---|---|---|---|---|---|---|---|---|
| *Spizellomyces punctatus* | Random-walk | 3-5[24,25] | Globular[24,25] | 20-24[24,25] | $1A_2$[24,26] | No[24,26] | Yes[24,26] | No[24,26] | [24,26] 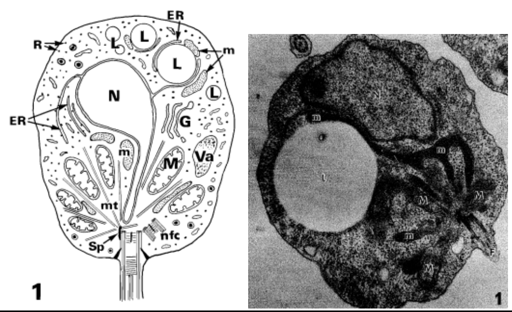 |
| *Geranomyces variabilis* | Random-walk | 3-5[27–29] | Globular[27–29] | 18.9-24.4[27–29] | $1A_2$[1,27,28] | No[27,28] | Yes[27,28] | No[27,28] | [27] 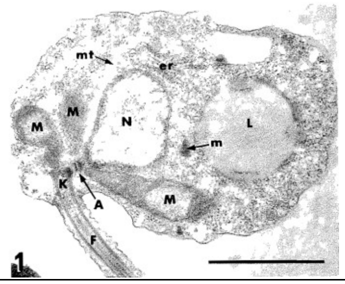 |





Table S1. Main cell biology characteristics of the zoospores of 12 species of Blastocladiomycota and Chytridiomycota, used to reconstruct Figure 4.
